# Longitudinal immune cell profiling in early systemic lupus erythematosus

**DOI:** 10.1101/2021.11.08.467791

**Authors:** Takanori Sasaki, Sabrina Bracero, Joshua Keegan, Lin Chen, Ye Cao, Emma Stevens, Yujie Qu, Guoxing Wang, Jennifer Nguyen, Stephen E. Alves, James A. Lederer, Karen H. Costenbader, Deepak A. Rao

## Abstract

**Objective:** To investigate the immune cell profiling and their longitudinal changes in systemic lupus erythematosus (SLE).

**Methods:** We employed mass cytometry with three different 38-39 marker panels (Immunophenotyping, T cell/monocyte, and B cell) in cryopreserved peripheral blood mononuclear cells (PBMCs) from nine patients with early SLE, 15 patients with established SLE, and 14 non-inflammatory controls. We used machine learning-driven clustering, FlowSOM (Flow Self-Organizing Maps) and dimensional reduction with tSNE (t-distributed Stochastic Neighbor Embedding) to identify unique cell populations in early and established SLE. For the nine early SLE patients, longitudinal mass cytometry analysis was applied to PBMCs at three time points (at enrollment, six months post-enrollment, and one year post-enrollment). Serum samples were also assayed for 65 cytokines by Luminex multiplex assay, and associations between cell types and cytokines/chemokines assessed.

**Results:** T peripheral helper cells (Tph cells), T follicular helper cells (Tfh cells) and several Ki67^+^ proliferating subsets (ICOS^+^ Ki67^+^ CD8 T cells, Ki67^+^ regulatory T cells, CD19^int^ Ki67^hi^ plasmablasts, and Ki67^hi^ PU.1^hi^ monocytes) were increased in early SLE. Longitudinal mass cytometry and multiplex serum cytokine assays of samples from early SLE patients revealed that Tfh cells and CXCL10 decreased at one year post-enrollment. CXCL13 correlated positively with several of the expanded cell populations in early SLE.

**Conclusions:** Two major helper T cell subsets and unique Ki67^+^ proliferating immune cell subsets were expanded in the early phase of SLE, and the immunologic features characteristic of early SLE evolved over time.

---

Systemic lupus erythematosus (SLE) is a prototypical autoimmune disease that affects multiple vital organs. Untreated immune activation in SLE can lead to tissue inflammation and irreversible organ damage, thus rapid recognition of lupus disease activity is an important goal in the care of patients with SLE. Delays in treatment are associated with poorer treatment responses and worse outcomes (1-3).

Despite the importance of early recognition and intervention, diagnosis of early SLE is often difficult because initial manifestations of the disease frequently include relatively non-specific symptoms. Fever, autoantibody production, hypocomplementemia and leukopenia are relatively common in early SLE (4), indicating that systemic immunologic features are already altered in the early phase. We hypothesize that defining the alterations in immune cell populations early in disease will provide critical insights into the early evolution of pathologic immune cell activation in SLE and may yield key metrics to diagnose SLE in the early phase.

A series of single-cell RNA sequencing studies from inflamed tissues recently identified aberrant immune cell expansions and cytokine/chemokine-mediated cellular networks within the affected organs in SLE (5, 6). These unbiased analyses provided broad and robust information on the composition of the immune cell infiltrates in the kidney in lupus nephritis. However, since multiple biopsies are difficult in most cases and most of the tissue samples were obtained from patients with established disease, there is limited information on immunologic features early in disease and little description of changes in inflammatory features between early and later established phases of disease. This information is important for the definition of immune response evolution over time in lupus, which may provide insights into differences in treatment response over time. From this perspective, blood samples are easier to access and analyse longitudinally, yet few longitudinal studies of associations between immunophenotype and clinical features in SLE over time have been reported (7, 8).

Mass cytometry (or Cytometry by Time Of Flight, CyTOF) is a powerful tool to broadly assess surface markers as well as intracellular proteins on immune cells. Dimensional reduction and visualization with tSNE (t-distributed Stochastic Neighbor Embedding) (9) combined with machine learning-driven clustering with methods such as FlowSOM (Flow Self-Organizing Maps) (10) allow for discrimination of distinct immune cell clusters in an unbiased way. Previously, the increase of PD-1^hi^ CXCR5^-^ CCR2^+^ CD4 T cells (T peripheral helper cells; Tph cells) in the peripheral blood of patients with SLE was identified by this methodology (11). CCR2 is a homing protein that promotes migration of immune cells to inflammatory sites, suggesting that the increase of the circulating immune cell subset reflects an inflammatory condition in the affected sites. Two studies have recently reported analyses using mass cytometry to examine blood samples from patients with established SLE (12, 13), but the longitudinal changes have not yet been studied.

Here we report broad mass cytometry data analyses with three different 38-39 marker panels in blood cells from patients with a new diagnosis of SLE. We first identified several unique immune cell populations in early SLE through unsupervised clustering and then verified the frequencies of these immune cell subsets. We further investigated the immune cell frequencies and serum cytokine levels in early SLE over time (at enrollment, six months, and 12 months post-enrollment). These longitudinal analyses indicated that several unique Ki67^+^ proliferating immune cell subsets are expanded even in the early phase of SLE and remain consistently elevated over time. In contrast, T follicular helper cells (Tfh cells) appeared elevated early after diagnosis and decreased over time. Serum cytokine profiling identified CXCL10, CD40L, IL-20, and TWEAK increased in early SLE, but among them, CXCL10 decreased longitudinally. Our data provide a detailed assessment of the immunologic features characteristic of early SLE as well as their changes over time.

## Results

### An unsupervised cell clustering view of the immune cell landscape in SLE

To investigate immunological and longitudinal changes in SLE, we evaluated cross-sectional and longitudinal analyses of peripheral blood mononuclear cells (PBMCs) by mass cytometry using three different panels (broad immunophenotype panel, T cell/monocyte-focused panel, and B cell-focused panel), along with 65-analyte serum cytokine profiling data (**Figure 1A, S-Table 1**). As an overview of our approach, we first applied unsupervised cell clustering using FlowSOM and dimensionality reduction by tSNE to the cross-sectional mass cytometry data from nine patients with early SLE who were enrolled within six months after the diagnosis, 15 patients with established SLE, and 14 non-inflammatory controls to identify distinct populations in an unbiased way. Early SLE patients were younger than established SLE patients (21.6 vs 36.5 years old, P<0.001), and corticosteroid (CS) use (33.3 vs 93.3%, P=0.03) and the dose (2.5 vs 14.5 mg/day, P=0.01) were higher in established SLE patients. SLEDAI-2K disease activity were comparable between the two groups (5.8 vs 5.1, P=0.80) (**S-Table 2, 3)**. We then investigated the longitudinal changes of the distinct immune cell populations and 65 cytokine levels from the nine patients with early SLE (**Figure 1B**), at three time points (A = at enrollment, B = six months after the enrollment, C = 12 months after the enrollment). Finally, we analysed associations between cell types and cytokines by hierarchical clustering.

**Figure 1.**
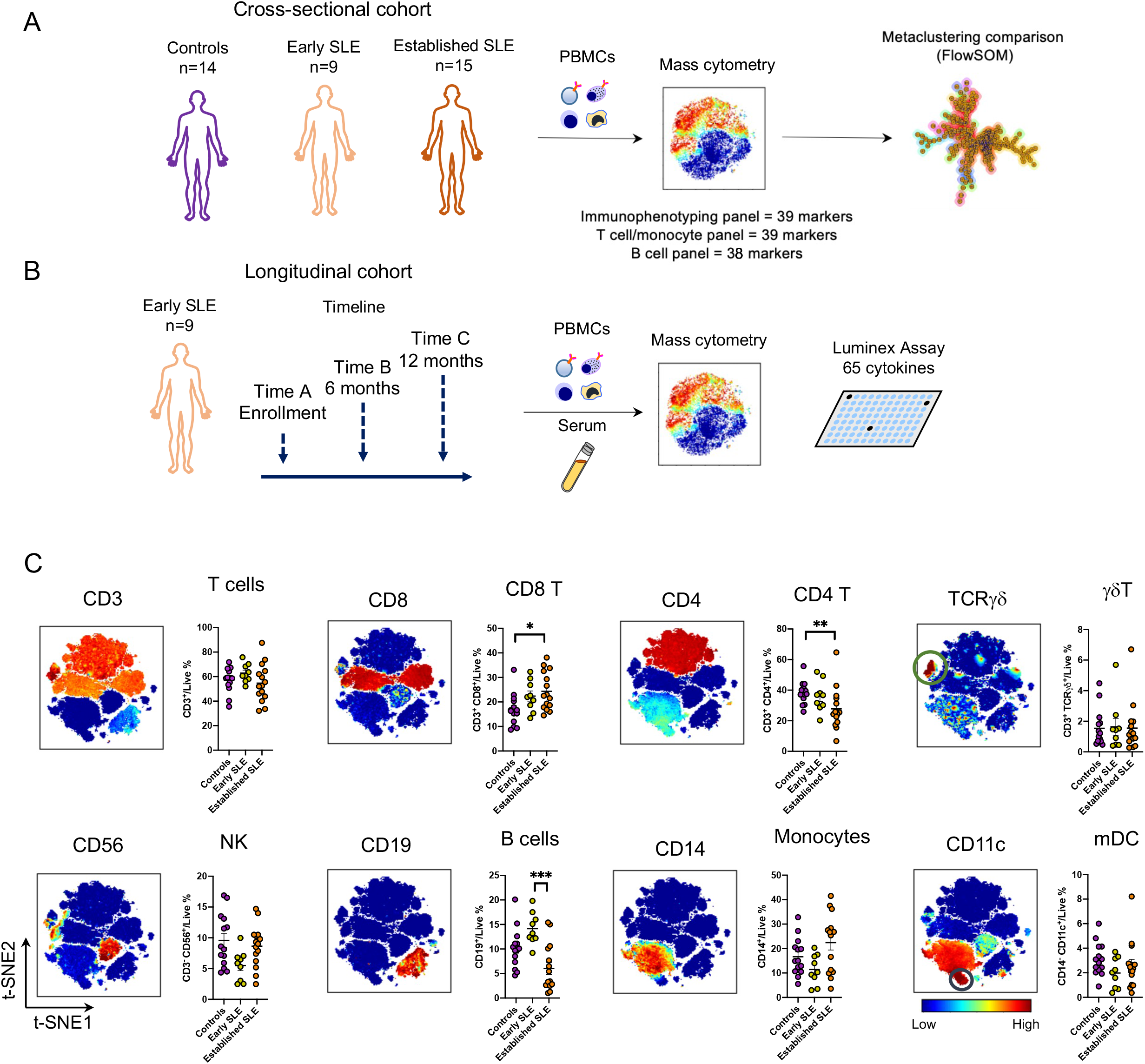
Overview of immunophenotyping early SLE using cross-sectional and longitudinal samples. (A) PBMCs were acquired from 14 controls, nine early SLE, and 15 established SLE and profiled with three different mass cytometry panels. (B) For the nine early SLE patients, longitudinal PBMC and serum samples were collected reflecting three timepoints: at enrollment (time A), six months after enrollment (time B), and 12 months after enrollment (time C). Immune cell frequency was assessed with mass cytometry and serum cytokines were assessed by Luminex cytokines assay. (C) Immunophenotyping panel mass cytometry data from nine early SLE and 15 established SLE were merged and visualized with tSNE plots. Frequencies of labelled immune cell types were compared between controls, early SLE, and established SLE cohorts by Kruskal-Wallis with Dunn’s test of multiple comparisons. *P<0.05, **P<0.01, ***P<0.001. Data are shown as mean ± SE.

For an initial, high-level view of the circulating immune cell populations, we first performed tSNE clustering of Immunophenotype panel mass cytometry data from the cross-sectional cohorts (**Figure 1C**). tSNE allowed clear visualization of distinct immune cell clusters, including three major CD3^+^ T cell populations, CD3^-^ CD56^+^ NK cells, CD19^+^ B cells, CD14^+^ monocytes, CD14^-^ CD11c^+^ myeloid dendritic cells (mDCs). Two of the CD3^+^ clusters were CD3^+^ CD4^+^ T cells and CD3^+^ CD8^+^ T cells. A third cluster, CD3^+^ CD4^-^ CD8^-^ T cells expressed TCR*γδ*, identifying this population as *γδ* T cells. Among these high-level immune cell subsets, the proportion of CD3^+^ CD8^+^ T cells was increased and the proportion of CD3^+^ CD4^+^ T cells was decreased in established SLE patients compared to controls, but none of these populations were higher in early SLE compared to controls.

### Expanded Ki67^+^ activated CD8 T cells in SLE patients

We next investigated changes in CD8 T cell populations in early SLE. To identify cell populations that differ between controls and SLE patients, we clustered CD8 T cells based on the 39-marker T cell-focused panel using FlowSOM. We compared the abundances of the clusters between SLE and controls and identified metacluster 10 as significantly increased in early SLE patients (5.2-fold, P<0.05, Kruskal-Wallis with Dunn’s multiple comparisons test) (**Figure 2A**). Heatmap expression analysis revealed that metacluster 10 contained cells with high expression of Ki67 and ICOS, suggesting that they are proliferating CD8 T cells (**Figure 2B**). Metaclusters 5 and 9 showed a similar expression pattern to metacluster 10 with high expression of Ki67 and ICOS and tended to be higher in SLE (**Figure 2A**). tSNE visualization of merged data from early SLE patients confirmed that the Ki67^+^ proliferative population expressed ICOS (**Figure 2C**). This population also expressed PD-1 and HLA-DR, suggesting that these are activated CD8 T cells. Interestingly, the Ki67^+^ CD8 T cells did not highly express granzyme B and granzyme K. Biaxial plots demonstrated that Ki67^+^ ICOS^+^ CD8 T cells significantly increased in early SLE compared to controls (0.8 vs 3.5%, P<0.01, Kruskal-Wallis with Dunn’s multiple comparisons test) (**Figure 2D**).

**Figure 2.**
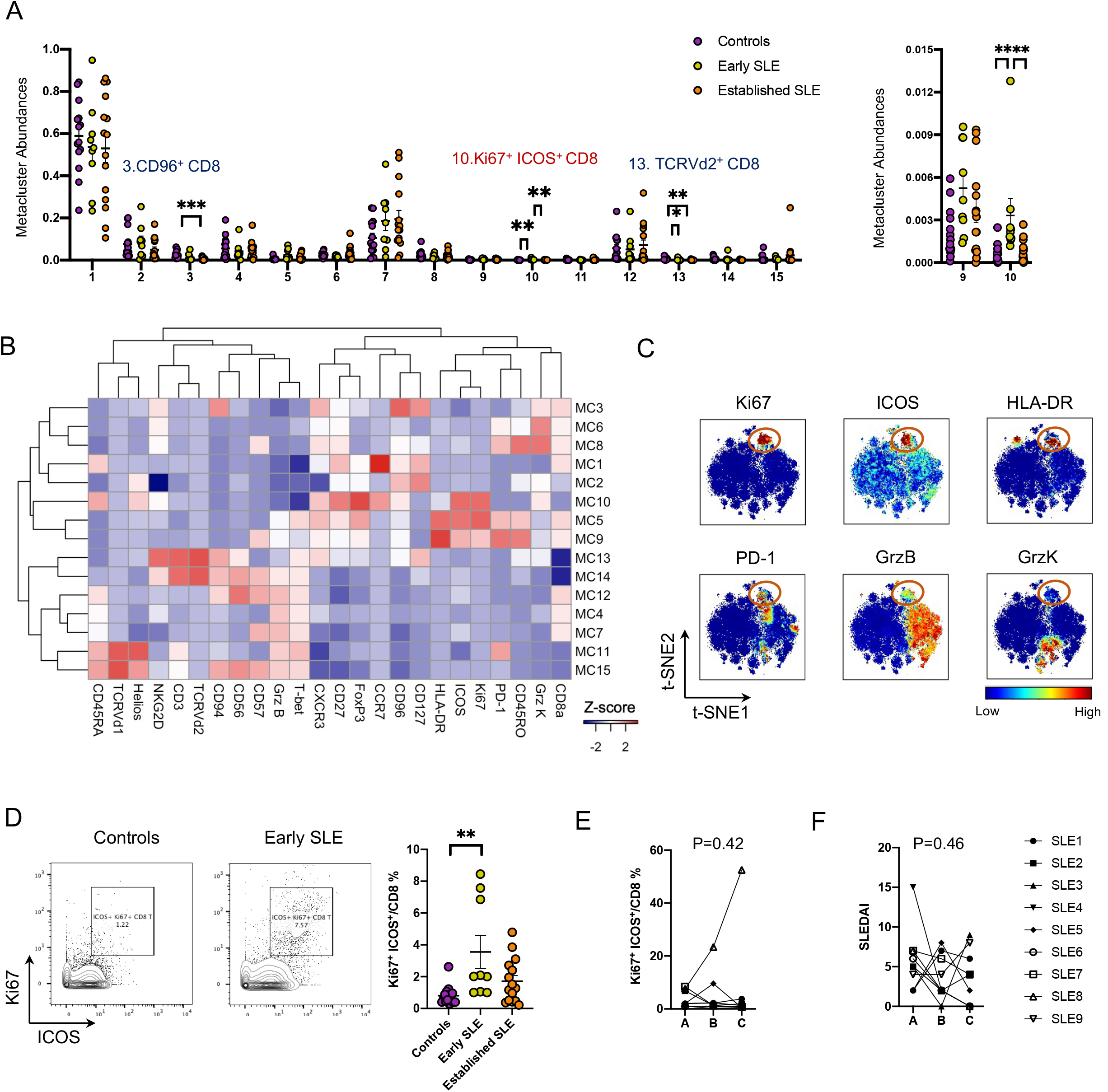
Expanded Ki67^+^ ICOS^+^ CD8 T cells in early SLE. (**A**) Abundance (% of total CD8 T cells) of FlowSOM metaclusters of CD8^+^ T cells in 14 controls, nine early SLE and 15 established SLE. Kruskal-Wallis with Dunn’s test was used for comparisons. (B) Heatmap of normalized expression of mass cytometry markers in each metacluster. Markers with average of metacluster medians > 0.2 are shown. (C) tSNE visualization of CD8 T cells from nine early SLE patients. The orange circle: Ki67^+^ ICOS^+^ CD8 cells. (D) Representative gating for Ki67^+^ ICOS^+^ CD8 T cell population in CD3^+^CD14^-^CD4^-^CD8^+^ cells, and the comparison between 14 controls, nine early SLE, and 15 established SLE. (E) Longitudinal changes of Ki67^+^ ICOS^+^ CD8 T cell frequency in early SLE at enrollment (time A), six months after enrollment (time B), and 12 months after enrollment (time C). Wilcoxon log-rank test to compare between A and C was used for p-value calculation. (F) Longitudinal changes of disease activity by SLEDAI in early SLE patients as in (E). *P<0.05, **P<0.01, ***P<0.001. Data are shown as mean ± SE.

In longitudinal analyses including time points six and 12 months post-enrollment, Ki67^+^ ICOS^+^ CD8 T cells remained persistently elevated over time. Disease activity in this patient cohort remained similarly active during this time frame (**Figure 2E, F**). We also identified metacluster 13, which contained T cells with high expression of CD94, CD56, and TCRV*δ*2, as significantly decreased in early SLE and established SLE patients (**Figure 2A, B**). Metacluster 3, which contained CD96^+^ CD8 T cells, was reduced in established SLE, but not in early SLE (**Figure 2A, B**).

### Tfh cells but not Tph cells decreased over time in early SLE

We next applied FlowSOM to CD4 T cells in controls and SLE patients. We identified metaclusters 6 (2.3-fold, P<0.01), 13 (5.8-fold, P<0.01), 14 (2.8-fold, P<0.05), and 15 (3.9-fold, P<0.01) significantly increased in early SLE patients at diagnosis (**Figure 3A**). Cells in metacluster 6 and metacluster 13 highly expressed PD-1, ICOS, and CD40L, but lacked CXCR5, suggesting that these two metaclusters contained Tph cells (**Figure 3B**). Interestingly, these two clusters were quite different in expression of CXCR3 and T-bet; low expression in metacluster 6 and high expression in metacluster 13. Metacluster 14 could be classified as Tfh cells with the high expression of PD-1, ICOS, and CXCR5. Metacluster 15 demonstrated a proliferating Treg phenotype with high expression of Ki67, FoxP3, CTLA-4, Helios, CD25, and CD39, and low expression of CD127 (**Figure 3B**). tSNE visualization revealed distinct clusters of PD-1 in either CXCR5^+^ or CXCR5^-^ regions and Ki67^+^ FoxP3-expressing Treg in early SLE patients (**Figure 3C**).

**Figure 3.**
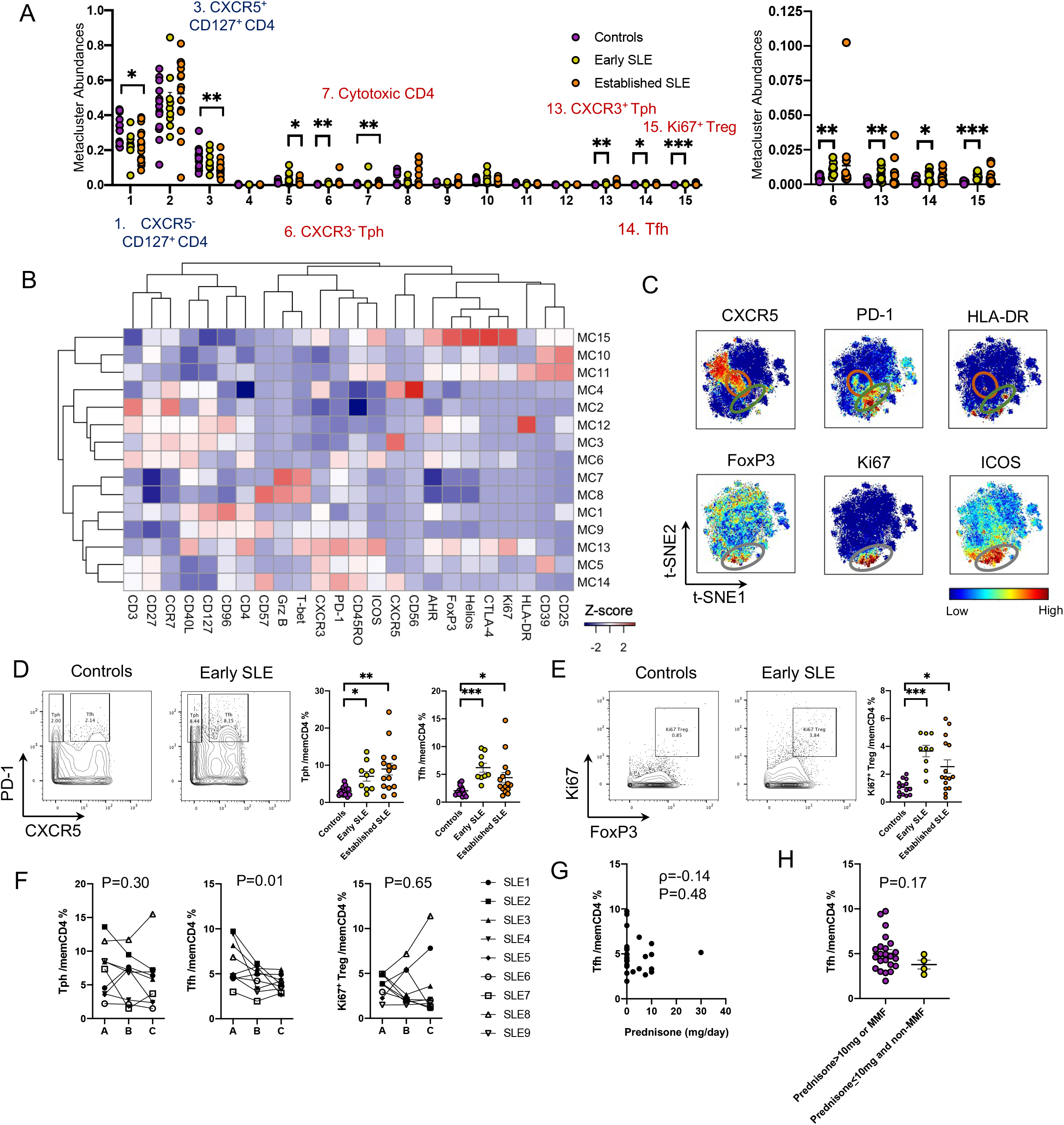
Distinct longitudinal changes of Tph cells and Tfh cells in early SLE. (A) Abundance (% of total CD4 T cells) of FlowSOM metaclusters of CD4^+^ T cells in 14 controls, nine early SLE and 15 established SLE. Kruskal-Wallis with Dunn’s multiple test was used for comparisons. (B) Heatmap of normalized expression of mass cytometry markers in each metacluster. Markers with average of metacluster medians > 0.2 are shown. (C) tSNE visualization of CD4 T cells in nine early SLE patients. The orange circle: Tfh cells, the green circle: Tph cells, the grey circle: Ki67^hi^ cells Treg cells. (D, E) Representative gating for Tph cells, Tfh cells, and Ki67^hi^ Treg in CD3^+^CD14^-^CD4^+^CD8^-^CD45RO^+^ memory CD4 T cells, and the comparison between 14 controls, nine early SLE, and 15 established SLE. Kruskal-Wallis with Dunn’s multiple test was used for comparisons. (F) Longitudinal changes of Tph cells, Tfh cells, and Ki67^+^ Treg cell in early SLE at enrollment (time A), six months after enrollment (time B), and 12 months after enrollment (time C). Wilcoxon log-rank test to compare between A and C was used for calculation of p-value. (G) Correlation between prednisone dose and Tfh cell frequency in 27 datapoints (nine early SLE patients x three timepoints each). Spearman statistics shown. (H) Tfh frequency in early SLE patients with or without MMF treatment as in (G) (n=27 data points). Mann-Whitney U test was used for the comparison. *P<0.05, **P<0.01, ***P<0.001. Data are shown as mean ± SE.

Quantification by biaxial gating confirmed that CXCR5^-^ PD-1^hi^ Tph cells (controls: 3.0%, early SLE: 7.0%, established SLE: 8.9%, P<0.05 in controls vs early SLE, P<0.01 in controls vs established SLE), CXCR5^+^ PD-1^hi^ Tfh cells (controls: 2.0%, early SLE: 6.2%, established SLE: 4.4%, P<0.001 in controls vs early SLE, P<0.05 in controls vs established SLE), and Ki67^+^ FoxP3^+^ Treg cells (controls: 1.0%, early SLE: 3.6%, established SLE: 2.5%, P<0.01 in controls vs early SLE, P<0.01 in controls vs established SLE) were increased in both early SLE and established SLE patients (**Figure 3D, E**). We also found that Tph cells and Ki67^+^ Treg cells were consistently elevated at one year after the diagnosis, whereas Tfh cells decreased over time in the early SLE cohort (**Figure 3F**). The proportion of Tfh cells did not correlate with corticosteroid dose and was not significantly different between the patients treated with prednisone >10mg and/or mycophenolate mofetil (MMF) and those without, suggesting that the decrease is independent from immunosuppressive treatment (**Figure 3G**). We identified metaclusters 1 and 3 significantly decreased in established SLE patients (**Figure 3A**). These metaclusters highly expressed CD127 and CD40L, but CXCR5 was quite distinct between metacluster 1 (CXCR5^-^) and metacluster 3 (CXCR5^+^) (**Figure 3B**).

### Increased CD19^int^ Ki67^hi^ plasmablasts in early SLE

We next applied the same clustering approach to CD19^+^ B cells, now using the B cell-focused mass cytometry panel (**S-Table 1**). We found metacluster 4 increased in early SLE patients (4.7-fold, P<0.05), metaclusters 11, 12, 13, and 15 increased in established SLE patients, and metacluster 14 increased in both early SLE (3.0-fold, P<0.05) and established SLE (**Figure 4A**). Expression heatmap analysis indicated that metacluster 4 contained a CD19^int^ Ki67^hi^ population with high expression of CD27 and CD38, indicating proliferating plasmablasts (**Figure 4B**). We also identified 5 metaclusters consistent with CD11c^+^ T-bet^+^ CD21^low^ CXCR5^-^ age-associated B cells (ABCs): HLA-DR^+^ CD38^-^ IgG^+^ ABCs (metacluster 11), HLA-DR^++^ CD38^+^ Ki67^+^ IgG^+^ ABCs (metacluster 12), CD1c^+^ IgM^+^ ABCs (metacluster 13), IgM^+^ IgD^+^ ABCs (metacluster 14), and CD11c^hi^ T-bet^hi^ ABCs (metacluster 15). tSNE visualization demonstrated distinct marker expression of CD19^int^ Ki67^hi^ plasmablasts and CD11c^+^ T-bet^+^ CD21^low^ CXCR5^-^ ABCs in early SLE (**Figure 4C**). Biaxial plots confirmed that the CD19^int^ Ki67^hi^ population contained CD27^+^ CD38^+^ plasmablasts, but CD19^hi^ Ki67^low^ population did not (**Figure 4D**). CD19^int^ Ki67^hi^ CD27^+^ CD38^+^ plasmablasts were significantly increased in early SLE patients (controls: 0.07%, early SLE: 0.88%, established SLE: 0.18%, P<0.01 in controls vs early SLE), whereas CD11c^+^ CD21^low^ ABCs were more expanded in established SLE patients (controls: 3.8%, early SLE: 7.6%, established SLE: 14.8%, P<0.001 in controls vs established SLE) (**Figure 4D, E**). CD19^int^ Ki67^hi^ CD27^+^ CD38^+^ plasmablasts were significantly lower in established SLE compared to early SLE patients, but also significantly lower in the SLE patients treated with prednisone >10mg and/or MMF compared the others (0.14% vs 1.1%, P<0.01) (**S-Figure 1**), suggesting that treatment may affect plasmablast abundance in treated SLE patients. Consistent with the increased abundance of metacluster 14 in early SLE, IgM^+^ IgD^+^ ABCs were significantly higher in early SLE patients compared to controls (**Figure 4F**). For other subclasses, almost 40% of CD19^int^ Ki67^hi^ CD27^+^ CD38^+^ plasmablasts were IgA, and IgG was rare, whereas IgG was more frequent (20%) in CD11c^+^ CD21^low^ ABCs (**Figure 4G**), indicating that class-switching isotypes were different in CD19^int^ Ki67^hi^ CD27^+^ CD38^+^ plasmablasts and CD11c^+^ CD21^low^ ABCs. In longitudinal analyses, CD19^int^ Ki67^hi^ CD27^+^ CD38^+^ plasmablasts and ABCs, and IgG or IgA class-switched CD19^int^ Ki67^hi^ CD27^+^ CD38^+^ plasmablasts and ABCs, stayed at high levels at one year after enrollment (**S-Figure 2, Figure 4H**). We also identified metacluster 6, which contained IgA^+^ IgD^-^ CD27^+^ memory B cells, as relatively decreased in early SLE patients.

**Figure 4.**
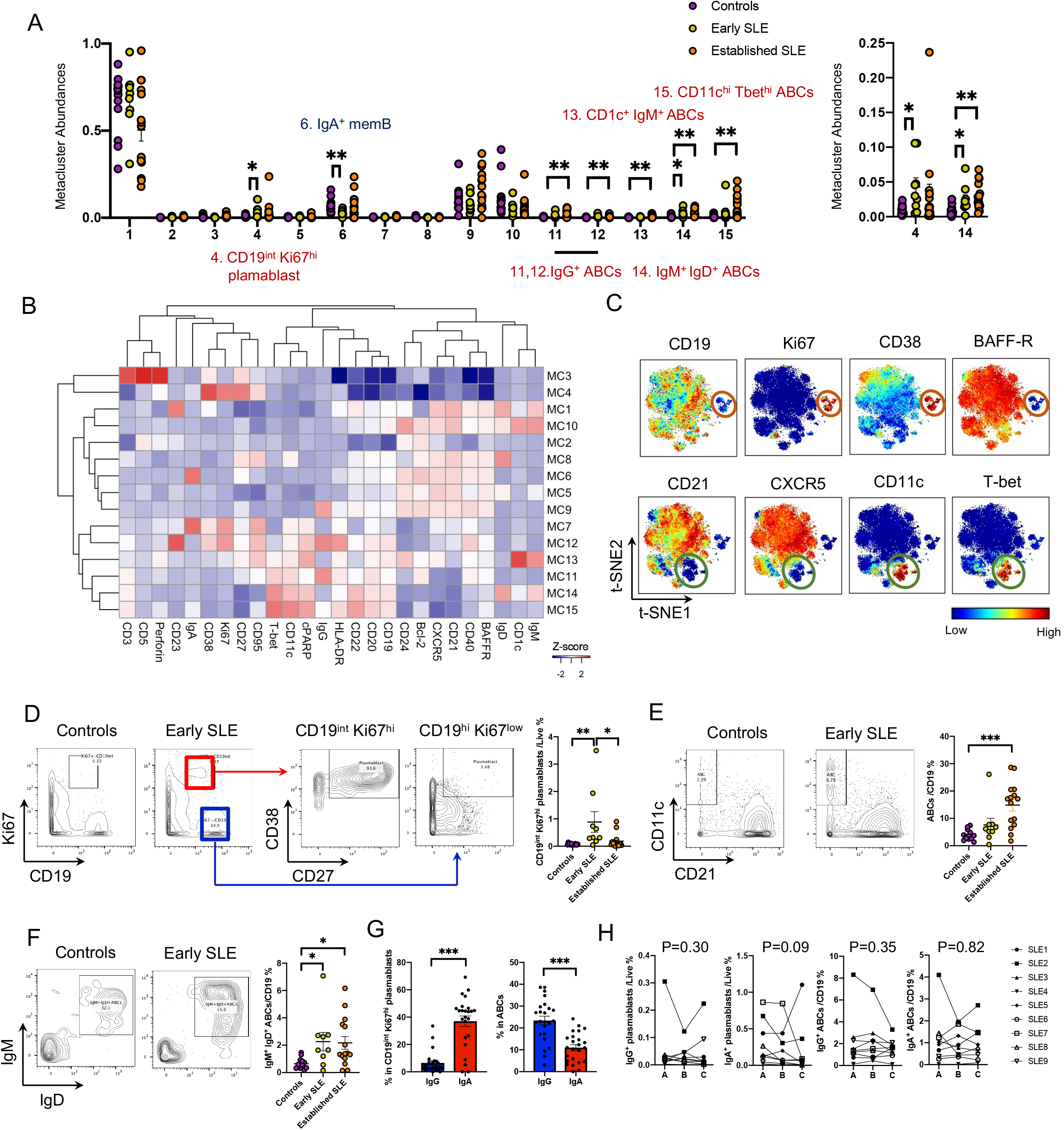
Expanded ABCs and plasmablasts in early SLE patients. (A) Abundance (% of total B cells) of FlowSOM metaclusters of B cells in 14 controls, nine early SLE and 15 established SLE. Kruskal-Wallis with Dunn’s multiple test was used for comparisons. (B) Heatmap of normalized expression of mass cytometry markers in each metacluster. Markers with average of metacluster medians > 0.2 are shown. (C) tSNE visualization of CD19^+^ B cells in nine early SLE patients. The orange circle: CD19^int^ Ki67^hi^ plasmablasts, the green circle: ABCs. (D-F) Representative gating for CD19^int^ Ki67^hi^ plasmablasts, ABCs, and IgM^+^ IgD^+^ ABCs in CD19^+^CD14^-^ B cells, and the comparison between 14 controls, nine early SLE, and 15 established SLE. Kruskal-Wallis with Dunn’s multiple test was used for comparisons. (G) Proportion of IgG^+^ and IgA^+^ cells in CD19^int^ Ki67^hi^ plasmablasts and ABCs in nine early SLE and 15 established SLE patients. Wilcoxon matched-pair signed rank test was used for the comparison. (H) Longitudinal changes of IgG^+^ CD19^int^ Ki67^hi^ plasmablasts, IgA^+^ CD19^int^ Ki67^hi^ plasmablasts, IgG^+^ ABCs, and IgA^+^ ABCs in early SLE at enrollment (time A), six months after enrollment (time B), and 12 months after enrollment (time C). Wilcoxon matched-pair signed rank test to compare between A and C was used for calculation of P value. *P<0.05, **P<0.01, ***P<0.001. Data are shown as mean ± SE.

### Increased PU.1^hi^ Ki67^hi^ monocytes in early SLE

In the CD14^+^ monocyte FlowSOM analysis using the T cell/monocyte panel, metacluster 8 was decreased and metacluster 13 was increased in early (5.6-fold, P<0.05) and established SLE patients compared to controls (**Figure 5A**). Heatmap analysis indicated that metacluster 13 contained HLA-DR^-^ PU.1^hi^ Ki67^hi^ monocytes (**Figure 5B**). tSNE visualization confirmed that Ki67^hi^ monocytes highly expressed PU.1 but not HLA-DR (**Figure 5C**). Biaxial plots revealed that metacluster 13 was well correlated with HLA-DR^-^ Ki67^hi^ monocytes and PU.1^hi^ Ki67^hi^ monocytes but more strongly in PU.1^hi^ Ki67^hi^ monocytes (**Figure 5D**). Consistent with the FlowSOM analysis, PU.1^hi^ Ki67^hi^ monocytes were increased in patients with early SLE, with comparable levels to established SLE patients (controls: 7.6%, early SLE: 29.6%, established SLE: 29.7%, P<0.05 in controls vs early SLE, P<0.05 in controls vs established SLE) (**Figure 5E**). PU.1^hi^ Ki67^hi^ monocytes expressed higher levels of CCR2 compared to PU.1^low^ Ki67^low^ monocytes (P<0.001) (**Figure 5F**). PU.1^hi^ Ki67^hi^ monocyte frequency did not change over time in early SLE patients (**Figire 5G**).

**Figure 5.**
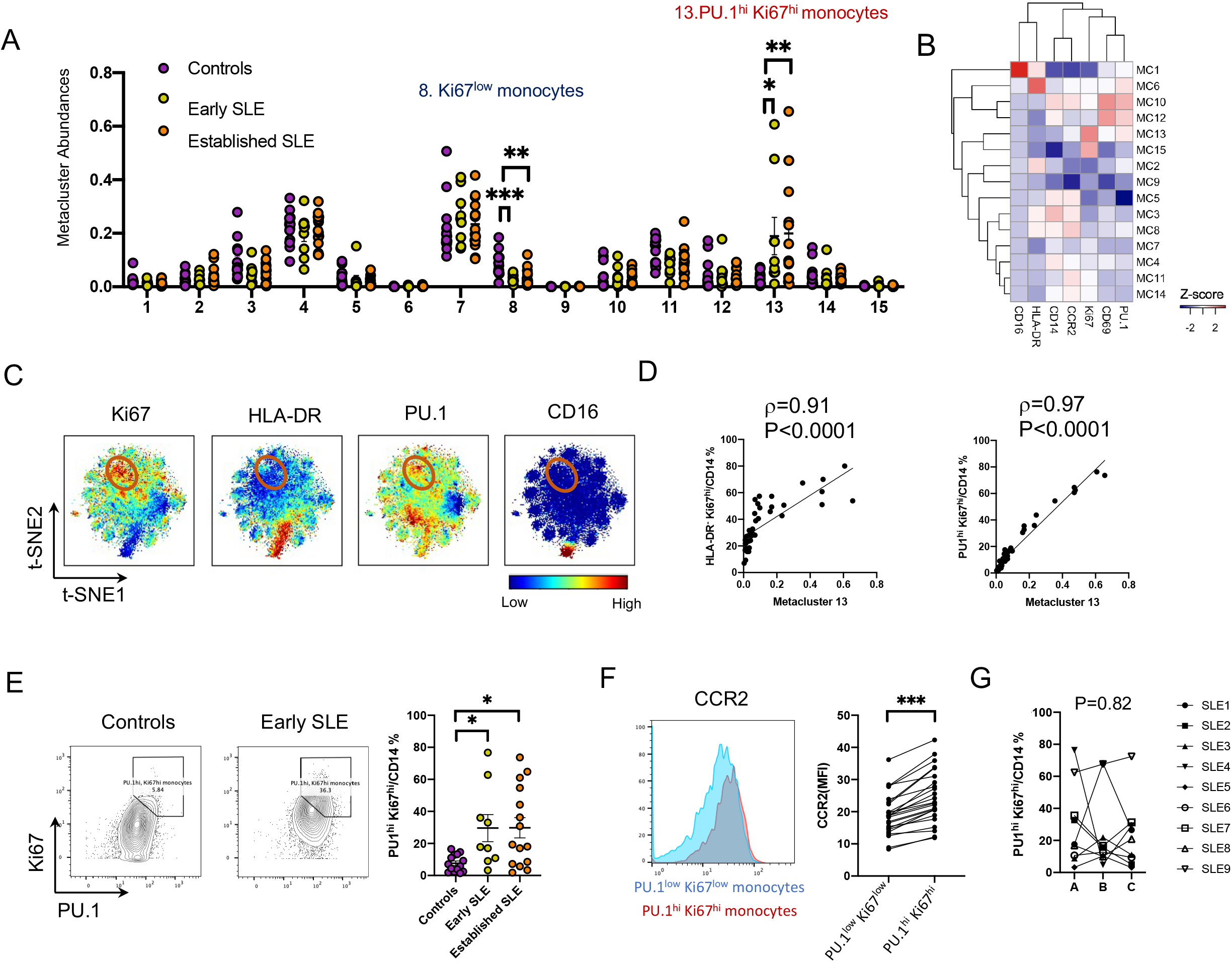
PU.1^hi^ Ki67^hi^ monocytes were expanded in early SLE. (A) Abundance (% of total monocytes) of FlowSOM metaclusters of monocytes in 14 controls, nine early SLE and 15 established SLE. Kruskal-Wallis with Dunn’s multiple test was used for comparisons. Kruskal-Wallis with Dunn’s multiple test was used for comparisons. (B) Heatmap of normalized expression of mass cytometry markers in each metacluster. Expression of PU.1, CD69, Ki67, CCR2, CD14, HLA-DR, CD16 are shown. (C) tSNE visualization of CD14 monocytes in nine early SLE. The orange circle: PU.1^hi^ Ki67^hi^ monocytes. (D) correlation of metacluster 13 with HLA-DR^-^ Ki67^hi^ monocytes and PU.1^hi^ Ki67^hi^ monocytes. Spearman correlation statistics shown. Solid line represents line of best fit. (E) Representative gating for PU.1^hi^ Ki67^hi^ monocytes and the comparison between 14 controls, nine early SLE, and 15 established SLE. Kruskal-Wallis with Dunn’s multiple test was used for comparisons. (F) CCR2 expression on PU.1^hi^ Ki67^hi^ monocytes and PU.1^low^ Ki67^low^ monocytes (nine early SLE and 15 established SLE). (G) Longitudinal changes of PU.1^hi^ Ki67^hi^ monocytes in 9 early SLE at enrollment (time A), six months after enrollment (time B), and 12 months after enrollment (time C). Wilcoxon log-rank test to compare between A and C was used for calculation of P value. *P<0.05, **P<0.01, ***P<0.001. Data are shown as mean ± SE.

### Associations between expanded immune cell populations in SLE

Since the analyses across multiple mass cytometry panels revealed that several Ki67^+^ proliferating populations were expanded in patients with SLE, we hypothesized that Ki67^+^ NK cells would also be increased in SLE. As we expected, biaxial plots indicated that NKG2D^+^ Ki67^+^ CD3^-^ CD56^+^ NK cells were highly increased in SLE, with stable levels in patients with early SLE over time (**Figure 6A, B**). The Ki67^+^ population did not express PD-1, HLA-DR, and ICOS, unlike Ki67^+^ CD4 or CD8 T cells (**Figure 6C**). Next, to identify associations between expanded immune cell populations in early SLE, we applied a hierarchical clustering analysis using the frequencies of Ki67^+^ ICOS^+^ CD8 T cells, Tph cells, Tfh cells, IgG^+^ CD19^int^ Ki67^hi^ plasmablasts, IgA^+^ CD19^int^ Ki67^hi^ plasmablasts, IgG^+^ ABCs, IgA^+^ ABCs, PU.1^hi^ Ki67^hi^ monocytes, and NKG2D^+^ Ki67^+^ NK cells. This analysis segregated cell populations into clusters with distinct patterns, including one cluster of PU.1^hi^ Ki67^hi^ monocytes and NKG2D^+^ Ki67^+^ NK cells (innate immunity cluster), one cluster of Ki67^+^ ICOS^+^ CD8 T cells, Tph cells, and Tfh cells (T cell cluster), and one larger cluster of IgG^+^ CD19^int^ Ki67^hi^ plasmablasts, IgA^+^ CD19^int^ Ki67^hi^ plasmablasts, IgG^+^ ABCs, and IgA^+^ ABCs (B cell cluster) (**Figure 6D**). Notably, Tph cells correlated with ABCs (ρ=0.51 P=0.006) and CD19^int^ Ki67^hi^ plasmablast (ρ=0.43 P=0.01), whereas Tfh cells did not (**Figure 6E**).

**Figure 6.**
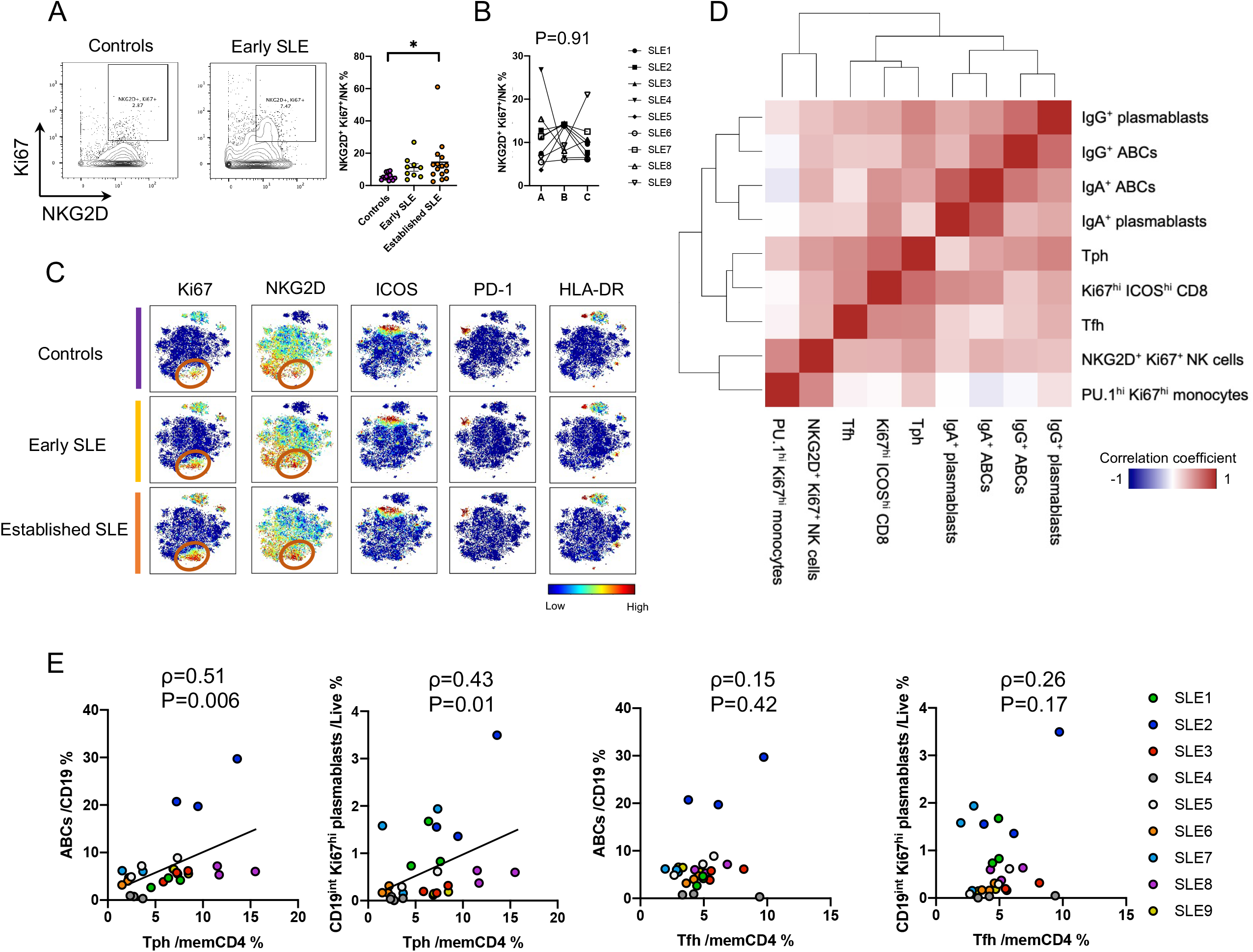
Association of Tph, Tfh, and Ki67 proliferative immune cells in early SLE. (A) Representative gating for NKG2D^+^ Ki67^+^ NK cells and the comparison between 14 controls, nine early SLE, and 15 established SLE. Kruskal-Wallis with Dunn’s multiple test was used for comparisons. (B) Longitudinal changes of NKG2D^+^ Ki67^+^ NK cells in nine early SLE patients at enrollment (time A), six months after enrollment (time B), and 12 months after enrollment (time C). Wilcoxon log-rank test to compare between A and C was used for calculation of P value. (C) tSNE visualization of NKG2D^+^ Ki67^+^ NK cells in 14 controls, nine early SLE, and 15 established SLE. The orange circle: NKG2D^+^ Ki67^+^ NK cells. (D) Hierarchical clustering heatmap with expanded immune cell types in early SLE. 27 data points from early SLE patients (nine patients x three time points) were used for this analysis. Correlation coefficients calculated by Spearman’s test were used for the heatmap. (E) Correlation analysis between Tph cells, Tfh cells, ABCs, and CD19^int^ Ki67^int^ plasmablasts. 27 early SLE patient data points were used as in (D). Spearman correlation statistics shown. Solid line represents line of best fit. *P<0.05, **P<0.01, ***P<0.001. Data are shown as mean ± SE.

### Longitudinal cytokine and chemokine profiling in early SLE

We next measured levels of 65 cytokines and chemokines in serum from 9 controls and nine early SLE patients, with the early SLE patients again analysed at three timepoints as in the cytometry analyses. Among the 65 cytokines/chemokines we selected *a priori* to be potentially important in early SLE pathogenesis, 33 cytokines were detected in serum samples. Interestingly, these cytokines positively correlated with each other together, suggesting a co-ordinately regulated underlying cytokine network in early SLE (**Figure 7A**). Most cytokines, with the exception of IL-16, had higher levels in early SLE samples, and IL-2R, CXCL10, CXCL13, IL-12p70, IL-17A, TSLP, CCL8, CCL24, TNF-RII, IL-2, IL-20, CD40L, CCL3, CD30, and TWEAK were significantly increased in early SLE samples, and CXCL10, CD40L, IL-20, and TWEAK remaining significantly higher even after Bonferroni correction to adjust for multiple testing (**Figure 7B, S-Figure 3, S-Figure 4**).Among these four cytokines, CXCL10 (P=0.03) was significantly decreased at 1 year after diagnosis, but CD40L, IL-20, and TWEAK stayed high levels. (**Figure 7C, S-Figure 5**). Next, to clarify the potential association between immune cells and chemokines, we investigated correlations between immune cell frequencies and serum chemokine levels in early SLE. Notably, CXCL13 broadly and strongly correlated with expanded lymphocyte subsets (Tph, Tfh, Ki67^hi^ ICOS^+^ CD8, ABC, plasmablasts) in samples from patients with SLE (**Figure 7D**). In contrast, CCL2 was strongly correlated with PU.1^hi^ Ki67^hi^ monocytes, the subset that highly expressed CCR2, suggesting the involvement of CCL2-CCR2 axis for PU.1^hi^ Ki67^hi^ monocytes migration to inflamed sites. These results suggest that different co-regulated pathways, which link cell types to related circulating factors, are active in early SLE patients.

**Figure 7.**
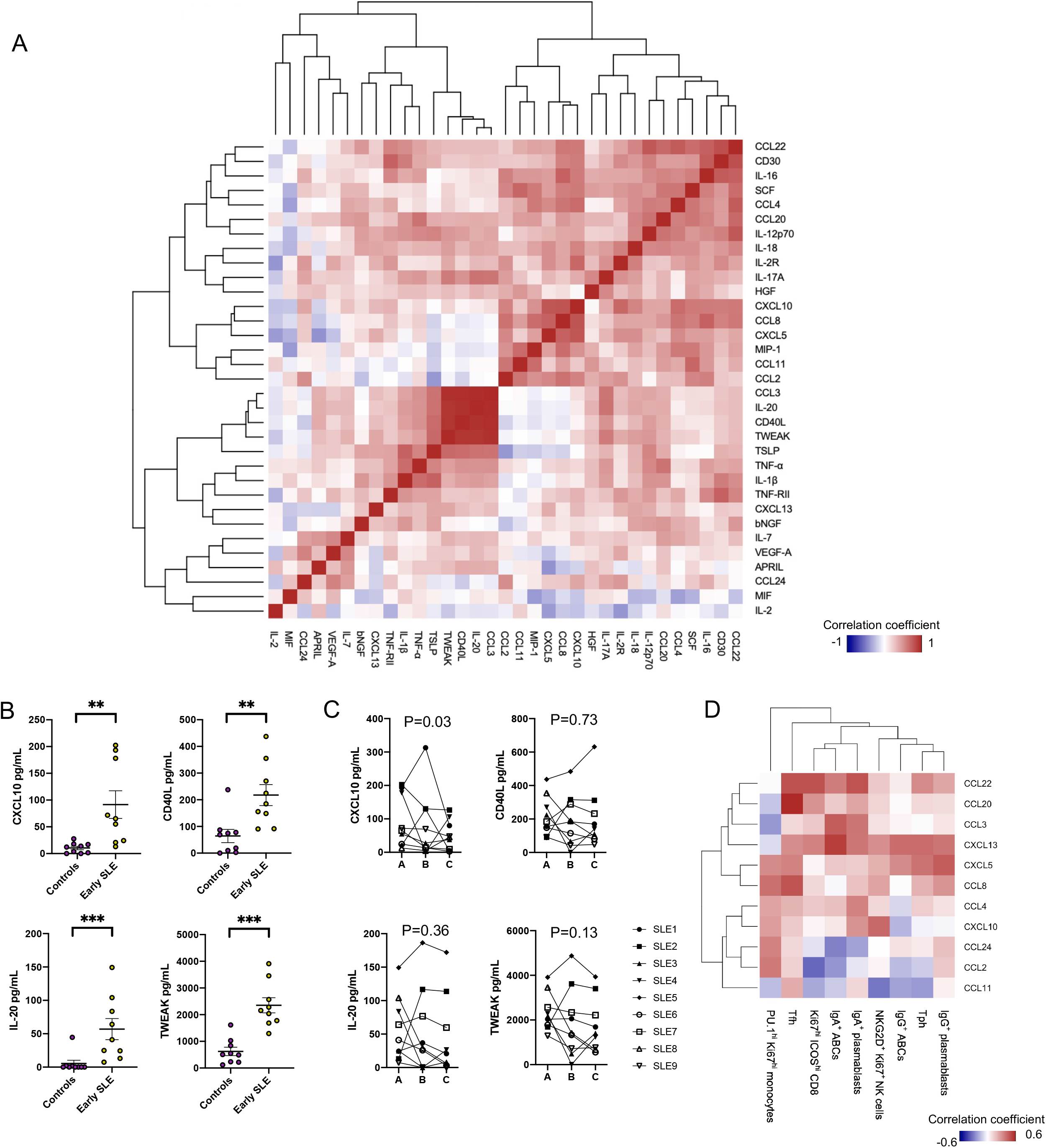
Dysregulated cytokine- and chemokine-networks in early SLE. (A) A hierarchical clustering heatmap with serum cytokines and chemokines in early SLE patients. 65 cytokines were assessed by Luminex assay, and 33 cytokines were detected. 27 data points from early SLE patients (nine patients x three time points) were used for this analysis. Correlation coefficients calculated by Spearman’s test were used for the heatmap. (B) Comparison of serum cytokine levels between nine controls and nine early SLE (time A). 15 cytokines were significantly different between controls and early SLE (time A). Mann-Whitney U test was used for the comparison. (C) Longitudinal changes of CXCL10, CD40L, IL-20, and TWEAK in nine early SLE at enrollment (time A), six months after enrollment (time B), and 12 months after enrollment (time C). Wilcoxon log-rank test to compare between A and C was used for calculation of p value. (D) A hierarchical clustering heatmap with expanded immune cell types and chemokines in early SLE. 27 data points from early SLE patients (nine patients x three time points) were used for this analysis. Correlation coefficients calculated by Spearman’s test were used for the heatmap. *P<0.05, **P<0.01, ***P<0.001. Data are shown as mean ± SE.

## Discussion

By broad and longitudinal cellular immunophenotyping and serum cytokine/chemokine profiling, we identified multiple expanded immune cell populations in patients with early SLE and evaluated their changes in the first year of disease and their relationships with serum cytokines/chemokines. We found that several lymphocyte populations expanded in early SLE share a common feature of expression of Ki67, a well-established marker of lymphocyte proliferation. This shared cytometric feature may capture the broad, active immune response occurring in early SLE. These Ki67^+^ populations, as well as Tph cells and ABCs, remain consistently elevated over the first year and are similarly elevated in established SLE patients, suggesting that these pathways are activated early and continue to characterize the pathologic immune response in SLE.

Notably, we also identified specific features of the immune response that change over time in early SLE patients. In particular, Tfh cells decreased over time in early SLE patients. Although Tfh cells and some of their inducing factors, such as IL-12, have been considered as therapeutic targets in SLE (14, 15), a phase III study of ustekinumab, a monoclonal antibody targeting interleukin IL-12 and IL-23, was discontinued due to the lack of efficacy (LOTUS study; NCT03517722). Since Tfh expanded initially, but decreased longitudinally, this target may have the therapeutic “window of opportunity”. Moreover, CD40L, IL-20, and TWEAK were increased at the initial time point and persistently elevated, while CXCL10 decreased over time. These data suggested that immune profiles change in each phase of SLE (**Figure 8**), such that quantification of some features of the immune response in SLE need to be adjusted based on disease duration.

**Figure 8.**
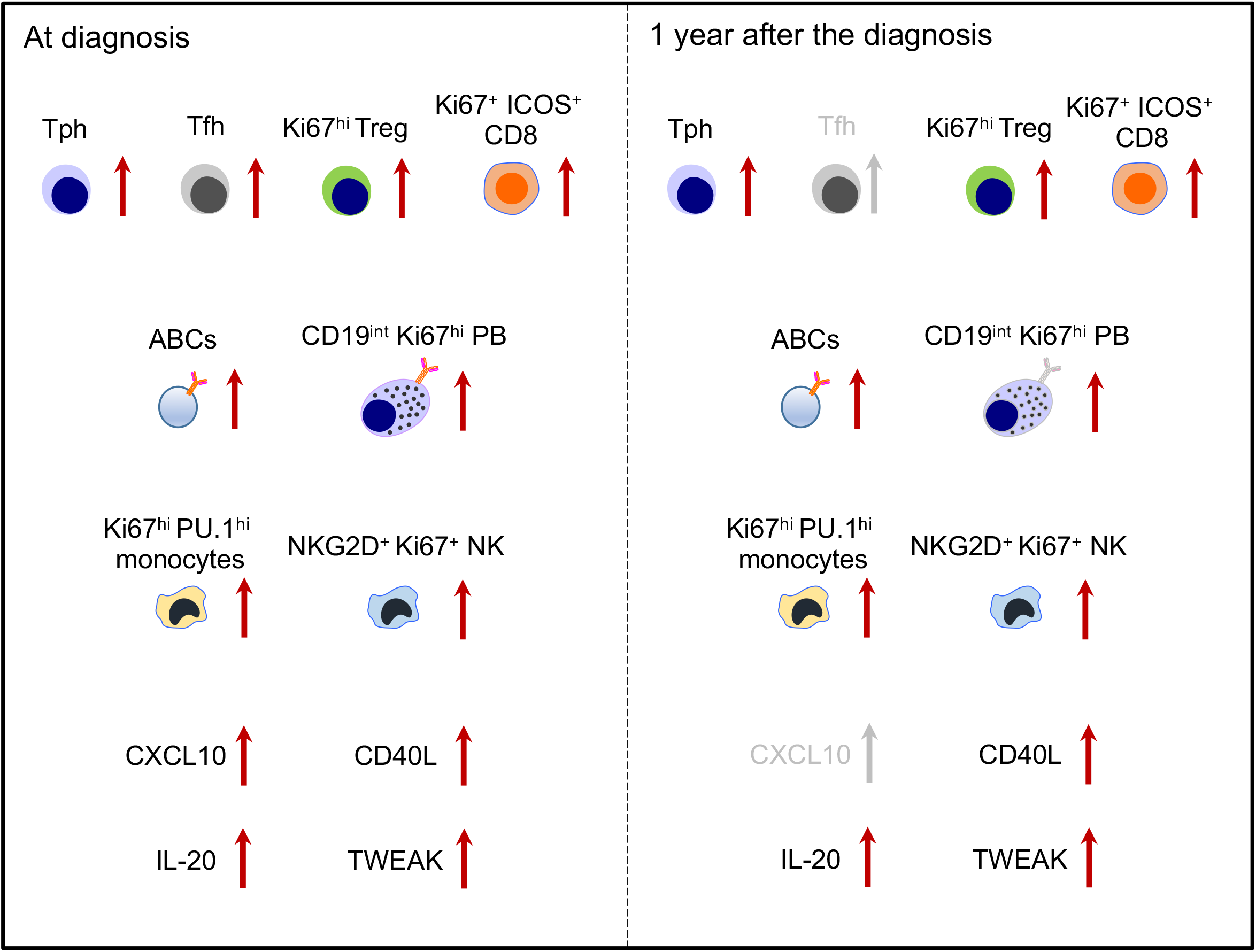
Dysregulated immune cell types, cytokines, chemokines, and chronological changes in early SLE. The increase pattern of cell types and cytokines/chemokines levels at the diagnosis and one year after the diagnosis in early SLE.

Diagnosing early SLE is challenging because the initial clinical manifestations are often non-specific. Identifying the immune system activation associated with early SLE may help to diagnose SLE as early as possible. Our study revealed that both antibody-secreting plasmablasts and B cell-helper T cells, including Tfh cells and Tph cells, were activated in the early phase and could be markers for early SLE. Since autoantibodies are increased several years prior to SLE onset (16), but not all autoantibody positive individuals develop SLE, it will be of major interest to determine whether alterations in circulating activated B cells and B cell-helper T can serve as specific hallmarks to predict risk of developing clinically evident SLE.

Both Tph cells and Tfh cells contribute to B cell responses through the production of IL-21, CD40L, and CXCL13 (17, 18). Strikingly, Tph cells stayed at high levels during the first year post-enrollment, whereas Tfh cells decreased longitudinally, suggesting distinct roles for Tph cells and Tfh cells over the course of the disease. One major difference between Tph cells and Tfh cells is their chemokine receptor expression, which determines their migratory capacity. Tph cells migrate into local inflammatory sites through receptors such as CCR2 and CCR5, while Tfh cells accumulate in B cell follicles within secondary lymphoid organs via a CXCR5-CXCL13 axis (19). Our data imply that Tfh-B cell interactions in secondary lymphoid organs may be particularly important at the initial onset of SLE, but the importance may shift to Tph-B cell interactions at local inflammatory sites over time. We did collect detailed data on SLE therapies administered to the newly diagnosed patients, but our small sample size precludes detailed analysis of how these therapies may impact lymphocyte populations and cytokines/chemokines over the first year of disease and this also deserves further study.

In the B cell analysis, the dominant subclasses differed in ABCs and CD19^int^ Ki67^hi^ plasmablasts. Recent broad BCR analysis of six different autoimmune diseases indicated that plasmablasts expressed more IgA1/2 compared to IgG1/2, whereas IgD^-^ CD27^-^ B cells, which contain much of the ABC population, expressed more IgG1/2 (20). Among these six autoimmune diseases, the frequency of IgA1/2 in PBMC B cells was higher in SLE, IgA vasculitis, Crohn’s disease, and Bechet’s disease compared to healthy controls. Considering that the mucosal associated lymphoid tissues (MALT) or gut-associated lymphoid tissues (GALT) are the main source of IgA^+^ plasmablasts (21), intestinal dysbiosis might be involved in the increase of plasmablasts in SLE.

We found that an expanded monocyte population in SLE expressed elevated levels of PU.1, a transcription factor implicated in macrophage development and function (22). Previous single-cell RNA-seq analyses of kidney biopsy samples suggested that inflammatory monocytes differentiate into phagocytic and M2-like macrophages in lupus nephritis (5). As PU.1^hi^ Ki67^hi^ monocytes expressed CCR2 and correlated well with serum CCL2, this monocyte population in the blood may be a precursor of inflammatory monocytes that infiltrate tissues.

A hierarchical clustering of immune cell subsets revealed distinct clusters of immune subsets with correlated abundance patterns, including clusters reflective of innate immunity, T cell activation, and B cell activation. Of note, Tph cells and Ki67^+^ ICOS^+^ CD8 T cells were strongly correlated, suggesting that these two subsets may be regulated through a common inducing factor. In this context, type I IFN may play an important role in the regulation. A series of RNA-seq analyses indicated that IFN signatures were highly enriched in Tph cells in SLE (11) and Ki67^hi^ CD8 T cells in immune checkpoint inhibitor-associated arthritis patients (23). In addition, type I IFN has negative regulatory effects on the expression of CXCR5 (24, 25). As Tph cells and Ki67^+^ ICOS^+^ CD8 T cells may be pathogenic drivers of SLE, anifrolumab, a fully human monoclonal antibody against the type I IFN receptor (26), may act to ameliorate the disease activity in part through the regulation of Tph cells and Ki67^+^ ICOS^+^ CD8 T cells.

Our study has several limitations. The relatively small cohort of early SLE patients followed limits the ability to identify co-correlated immune features and precludes evaluation of clinical correlates of the cellular features identified. A larger cohort will be required in subsequent studies to determine the potential prognostic significance of the immune features detected here. We have quantified cytometric and serum protein features but have not interrogated transcriptional programs. In addition, our study focuses only on blood samples and does not contain parallel tissue studies. Nonetheless, the substantial alterations demonstrated in circulating immune cells from patients with lupus support the idea that clinically relevant signals may be detectable in blood samples.

In conclusion, this study highlighted persistent activation of Tph, ABCs, and Ki67^+^ proliferating immune cells populations in the blood in early SLE and underscores the value of broad, longitudinal immunophenotyping to define patterns of SLE immune activity that may help refine potential biomarkers and prioritize therapeutic targets for early and established phases of SLE.

## Methods

### Study Subjects

All SLE patients met 1997 ACR classification criteria (27). For the early SLE cohort, nine SLE patient were who were within six months of disease diagnosis and without treatment with major immunosuppressive therapies (treatment with prednisone ≤ 10mg and hydroxychloroquine were permitted). For the cross-sectional study, 14 non-inflammatory controls and 15 patients with established SLE were also included. For the longitudinal cytometry study, the same nine patients with early SLE were evaluated at six months and 12 months after enrollment. For serum analyses, the same nine early SLE patients were evaluated, along with nine non-inflammatory controls different from the cross-sectional study. Detailed clinical information is shown in Supplementary Table 2, 3.

### Mass cytometry

Blood samples were collected into heparin tubes and PBMCs were isolated by density centrifugation using Ficoll-Hypaque in 50mL conical tubes. PBMCs were washed by PBS and cryopreserved in a 10% DMSO + 90% FBS solution. Samples from the cross-sectional cohorts as well as longitudinal samples from the early SLE cohort were collected and thawed together in batches of 20 samples per batch (total three batches) and processed for mass cytometry within a one-week period. The three longitudinal samples from each early SLE patient were included in the same batch to minimize potential batch effects.

Cryopreserved PBMCs were thawed into RPMI Medium 1640 (Life Technologies #11875-085) supplemented with 5% heat-inactivated fetal bovine serum (Life Technologies #16000044), 1 mM GlutaMAX (Life Technologies #35050079), antibiotic-antimycotic (Life Technologies #15240062), 2 mM MEM non-essential amino acids (Life Technologies #11140050), 10 mM HEPES (Life Technologies #15630080), 2.5 × 10^−5^ M 2-mercaptoethanol (Sigma-Aldrich #M3148), 20 units/mL sodium heparin (Sigma-Aldrich #H3393), and 25 units/mL benzonase nuclease (Sigma-Aldrich #E1014). Cells were counted and 0.5-1×10^6^ cells from each sample were transferred to a polypropylene plate for staining. The samples were spun down and aspirated. 5 μM of cisplatin viability staining reagent (Fluidigm #201064) was added for two minutes and then diluted with culture media. After centrifugation, Human TruStain FcX Fc receptor blocking reagent (BioLegend #422302) was used at a 1:100 dilution in PBS with 2.5 g bovine serum albumin (Sigma Aldrich #A3059) and 100 mg of sodium azide (Sigma Aldrich #71289) for 10 minutes followed by incubation with conjugated surface antibodies for 30 minutes. All antibodies were obtained from the Harvard Medical Area CyTOF Antibody Resource and Core (Boston, MA).

16% stock paraformaldehyde (Fisher Scientific #O4042-500) dissolved in PBS was used at a final concentration of 4% formaldehyde for 10 minutes in order to fix the samples before permeabilization with the FoxP3/Transcription Factor Staining Buffer Set (ThermoFisher Scientific #00-5523-00). The samples were incubated with SCN-EDTA coupled palladium based barcoding reagents for 15 minutes and then combined into a single sample. Conjugated intracellular antibodies were added into each tube and incubated for 30 minutes. Cells were then fixed with 1.6% formaldehyde for 10 minutes. DNA was labelled for 20 minutes with an 18.75 μM iridium intercalator solution (Fluidigm #201192B). Samples were subsequently washed and reconstituted in Milli-Q filtered distilled water in the presence of EQ Four Element Calibration beads (Fluidigm #201078) at a final concentration of 1×10^6^ cells/mL. Samples were acquired on a Helios CyTOF Mass Cytometer (Fluidigm).

### FlowSOM and tSNE analyses

The raw FCS files were normalized to reduce signal deviation between samples over the course of multi-day batch acquisitions, utilizing the bead standard normalization method established by Finck et al (28). These normalized files were then deconvoluted into individual sample files using a single-cell based debarcoding algorithm established by Zunder et al (29) Mass cytometry data were gated to exclude debris and identify DNA^+^ events. Non-viable cisplatin^+^ cells and equalization beads were excluded. FlowSOM analyses were performed using Cytobank. HCs (n=14) and SLE (n=24: early SLE n=9, established SLE n=15) samples were included. Metacluster and cluster numbers were 15 and 225, respectively. Each metacluster abundances were compared between HCs, early SLE, and established SLE. Heatmap analyses were performed with Z score of metacluster medians in each marker. For tSNE clustering, HCs and SLE fcs files were concatenated using FlowJo 10.4.2. tSNE analyses using Cytobank was performed with equal number events from the concatenated HCs or SLE fcs files. Gating for cell frequencies and expression intensity quantification were performed using FlowJo 10.4.2.

### Cytokines

65 cytokines (Supplementary figure 1) were measured by Luminex multiplex assay according to the manufacturer’s instructions. Associations between cytokines, and association cytokines and cell types assessed with heatmap analysis using Spearman’s correlation coefficient.

### Study approval

SLE patients and non-inflammatory controls were enrolled at Brigham and Women’s Hospital with informed consent under IRB protocols (2014P002558 and 2016P001660) approved by Mass General Brigham IRB.

### Statistics

Statistical comparisons were performed in Prism. Mann-Whitney test was used for comparison between two groups, and Kruskal-Wallis with Dunn’s multiple comparisons test for comparisons between three groups. Wilcoxon test was used for comparison between time point A and C in the longitudinal analyses. Correlation analysis was performed by Spearman’s test, and the best fit line were drawn when significant. Heatmap analysis was performed P-value < 0.05 (two sided) was regarded as significant. Data are shown as mean ± SE.

## Data availability

Data are available upon reasonable request to the corresponding author.

## Acknowledgements

This work is supported in part by funding from Merck Sharp & Dohme Corp., a subsidiary of Merck & Co., Inc., Kenilworth, NJ, USA, and NIAMS K24 AR066109 to KHC and from the Lupus Research Alliance, Burroughs Wellcome Fund Career Award in Medical Sciences, Doris Duke Charitable Foundation Clinical Scientist Development Award, NIAMS K08 AR072791 to DAR, and NIAMS P30 AR070253 to JAL and DAR.

