## Supplementary tables for "Longitudinal immune cell profiling in early systemic lupus erythematosus"

**Supplementary Table 1. Mass cytometry and cytokine panels in this study.**

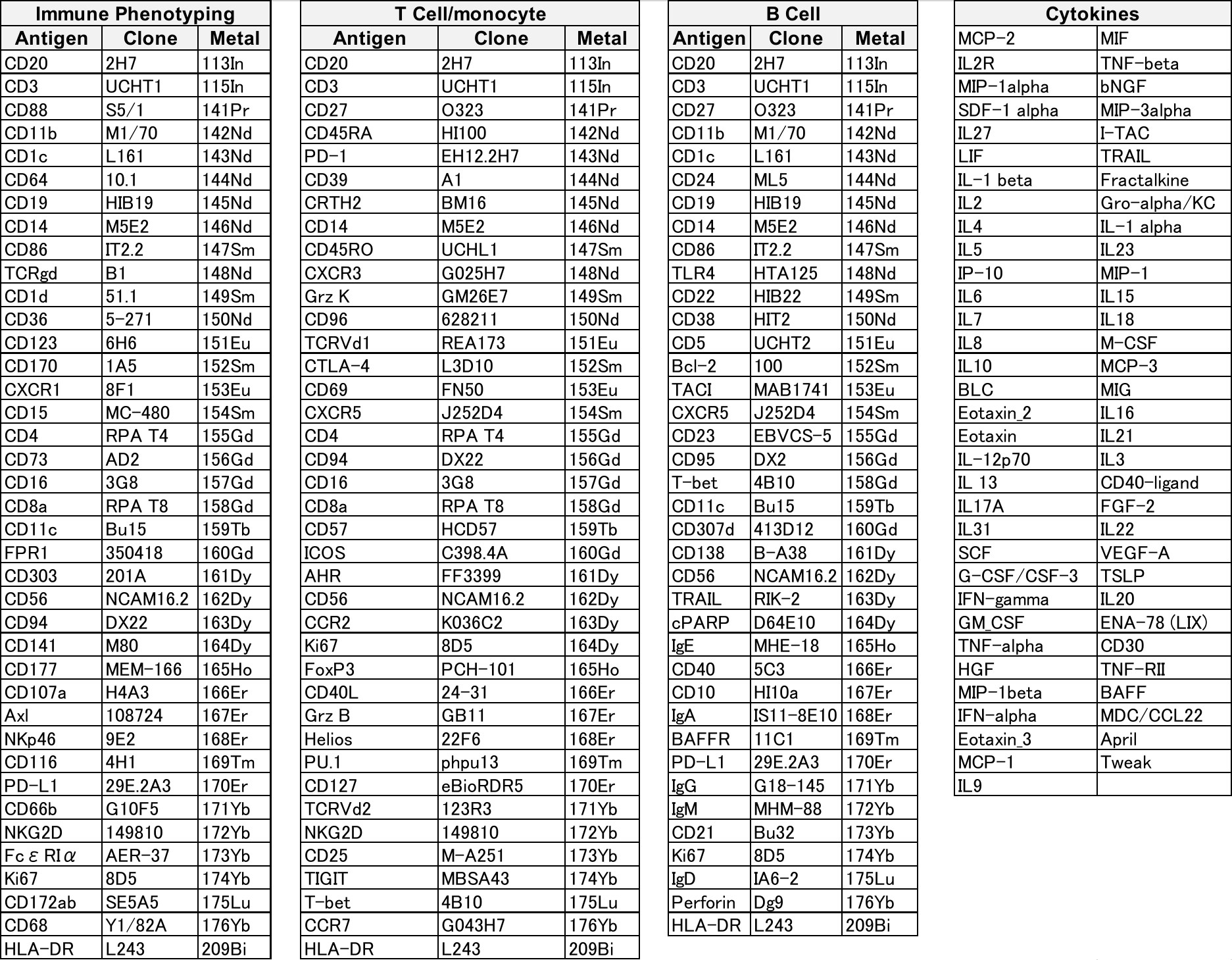

|  | **Controls (n=14)** | **Early SLE (n= 9)** | **Established SLE (n=15)** | **P value** |
| --- | --- | --- | --- | --- |
| Age | 46.1 ± 3.9 | 26.1 ± 1.6 | 36.5 ± 2.2 | <0.001 |
| Female N (%) | 10 (71.4) | 9 (100.0) | 14 (93.3) | 0.99 |
| SLEDAI 2k (total score) | N/A | 5.8 ± 1.2 | 5.4 ± 1.1 | 0.80 |
| Anti-dsDNA titer (IU/mL) | N/A | 168.2 ± 73.4 | 150.1 ± 50.9 | 0.82 |
| C3 (mg/dL) | N/A | 93.7 ± 13.2 | 97.2 ± 7.9 | 0.49 |
| C4 (mg/dL) | N/A | 16.2 ± 6.8 | 24.3 ± 6.8 | 0.35 |
| Mycophenolate  in past 6 months N (%) | N/A | 0 | 7 (46.6) | 0.02 |
| Cyclophosphamide  in past 6 months N (%) | N/A | 0 | 0 | 0.99 |
| Hydroxychloroquine  in past 6 months N (%) | N/A | 6 (66.6) | 11 (73.3) | 0.99 |
| Azathioprine  in past 6 months N (%) | N/A | 0 | 1 (6.6) | 0.99 |
| Corticosteriod use N (%) | N/A | 3 (33.3) | 14 (93.3) | 0.03 |
| Corticosteroid dose (mg/day) | N/A | 2.5 ± 0.8 | 14.5 ± 4.2 | 0.01 |

**Supplementary Table 2. Patient demographics in this study.**

Mann-Whitney U test or Fisher’s exact test were used for the comparison between early SLE and established SLE. Continuous variables are shown in mean ± SE.

**Supplementary Table 3. Longitudinal changes of disease activity and treatment in early SLE.**

Wilcoxon log-rank test or Fisher’s exact test were used for the comparison between early SLE and established SLE. Continuous variables are shown in mean ± SE.

|  | **At enrollment: Time A (n = 9)** | **At 6 months: Time B**  **(n = 9)** | **At 1 year: Time C (n = 9)** | **P value (A vs B)** | **P value (A vs C)** |
| --- | --- | --- | --- | --- | --- |
| SLEDAI 2k (total score) | 5.8 ± 1.2 | 3.6 ± 0.9 | 3.6 ± 1.2 | 0.45 | 0.46 |
| Mycophenolate use N (%) | 0 | 1 (11.1) | 2 (22.2) | 0.99 | 0.47 |
| Cyclophosphamide use N (%) | 0 | 0 | 0 | 0.99 | 0.99 |
| Hydroxychloroquine use N (%) | 6 (66.6) | 7 (77.7) | 8 (88.8) | 0.99 | 0.57 |
| Azathioprine use N (%) | 0 | 0 | 0 | 0.99 | 0.99 |
| Corticosteroid use N (%) | 3 (33.3) | 4 (44.4) | 3 (33.3) | 0.99 | 0.99 |
| Corticosteroid dose (mg/day) | 2.5 ± 0.8 | 5.8 ± 5.5 | 2.5 ± 1.3 | 0.50 | 0.99 |
