## Supplementary figures for "Longitudinal immune cell profiling in early systemic lupus erythematosus"

Supplementary figure 1

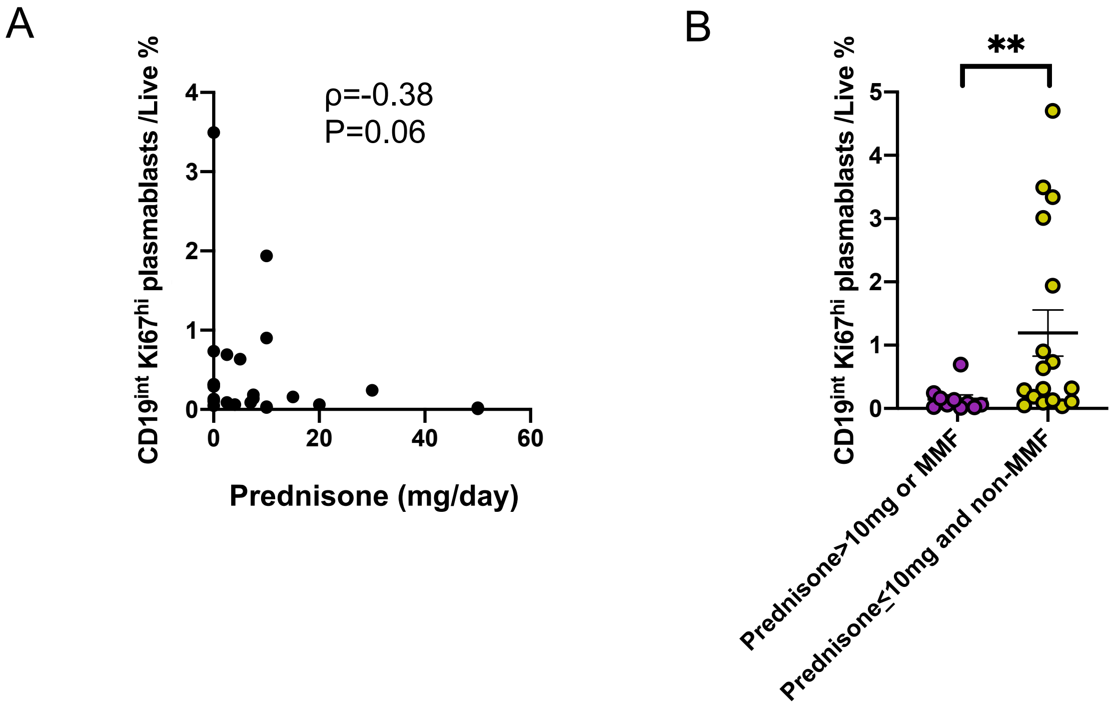

### Supplementary figure 2

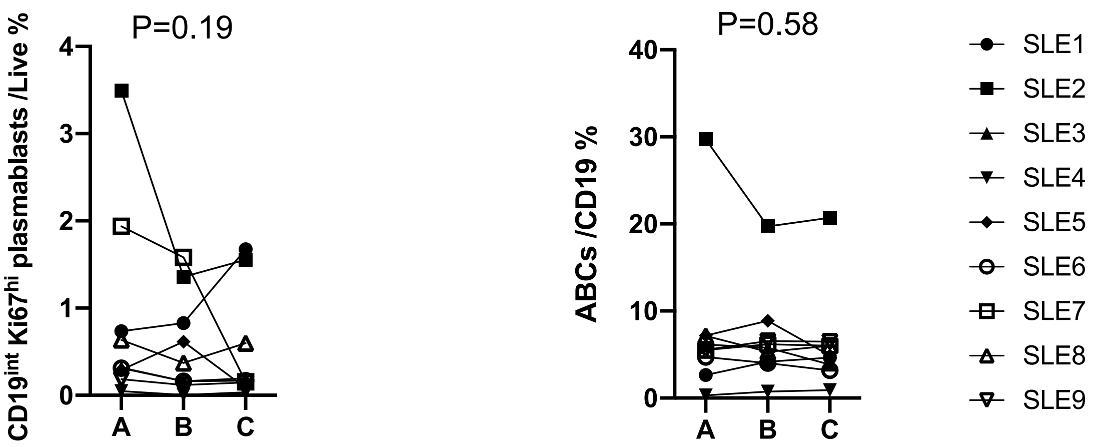

### Supplementary figure 3

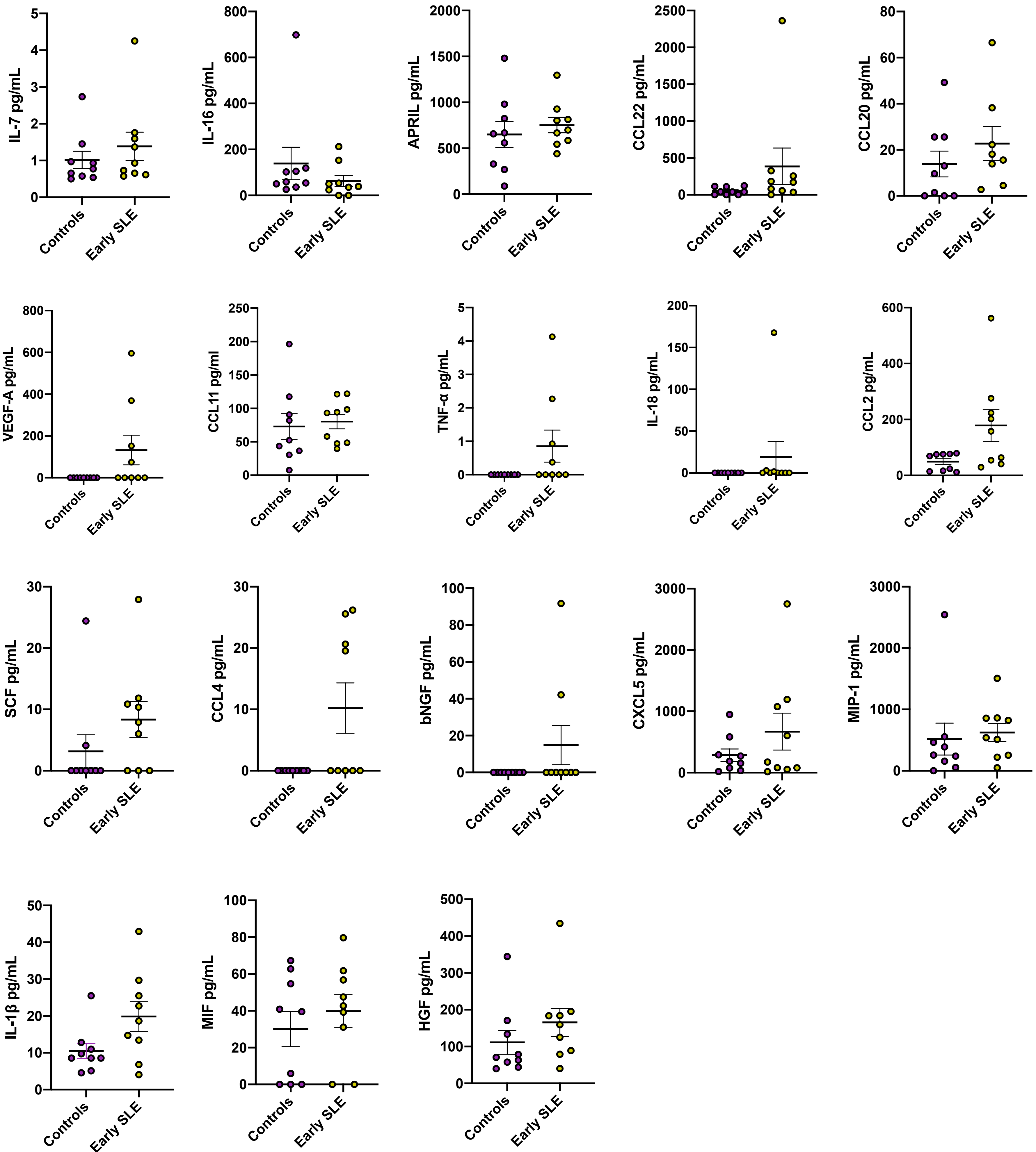

### Supplementary figure 4

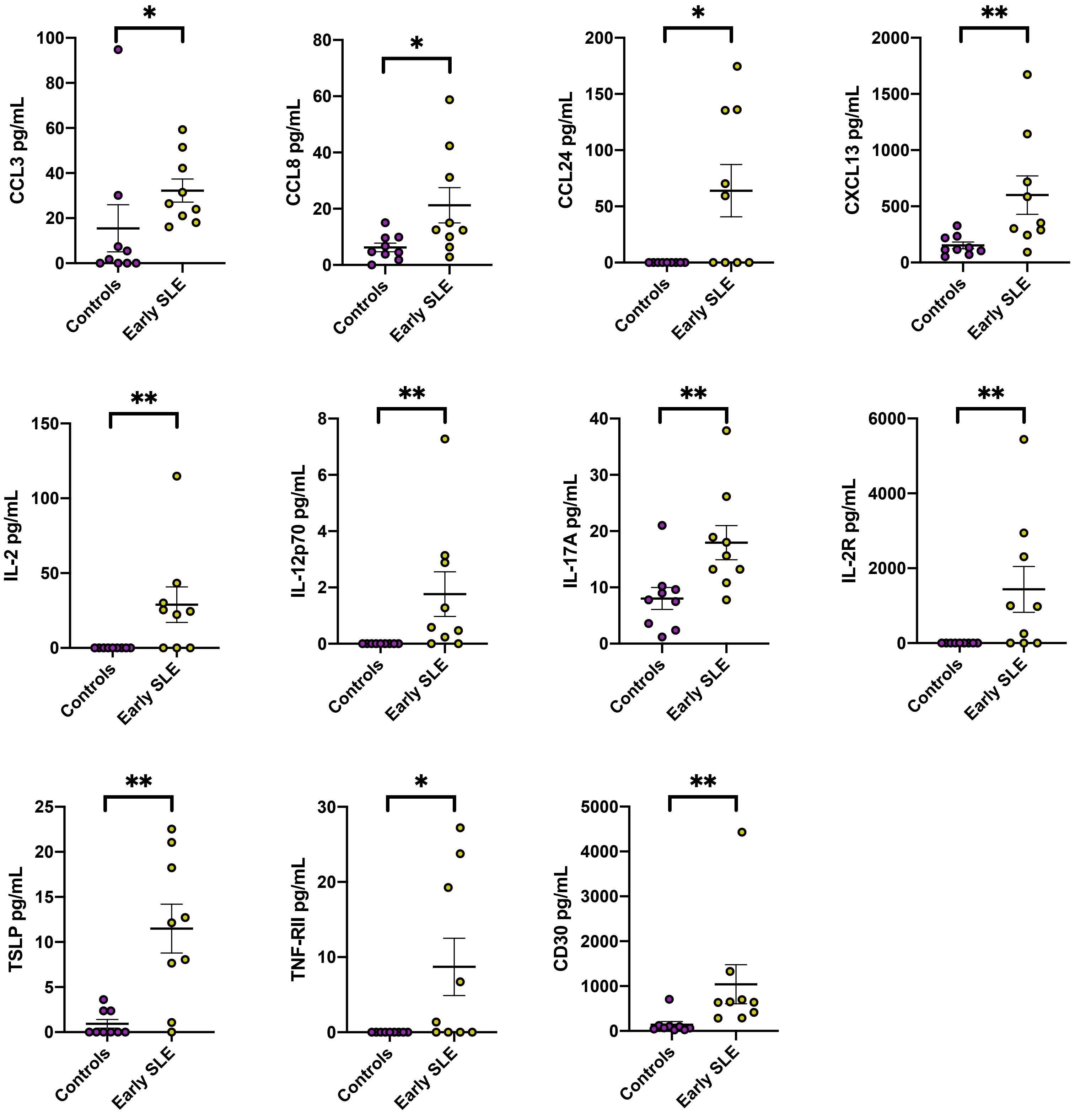

### Supplementary figure 5

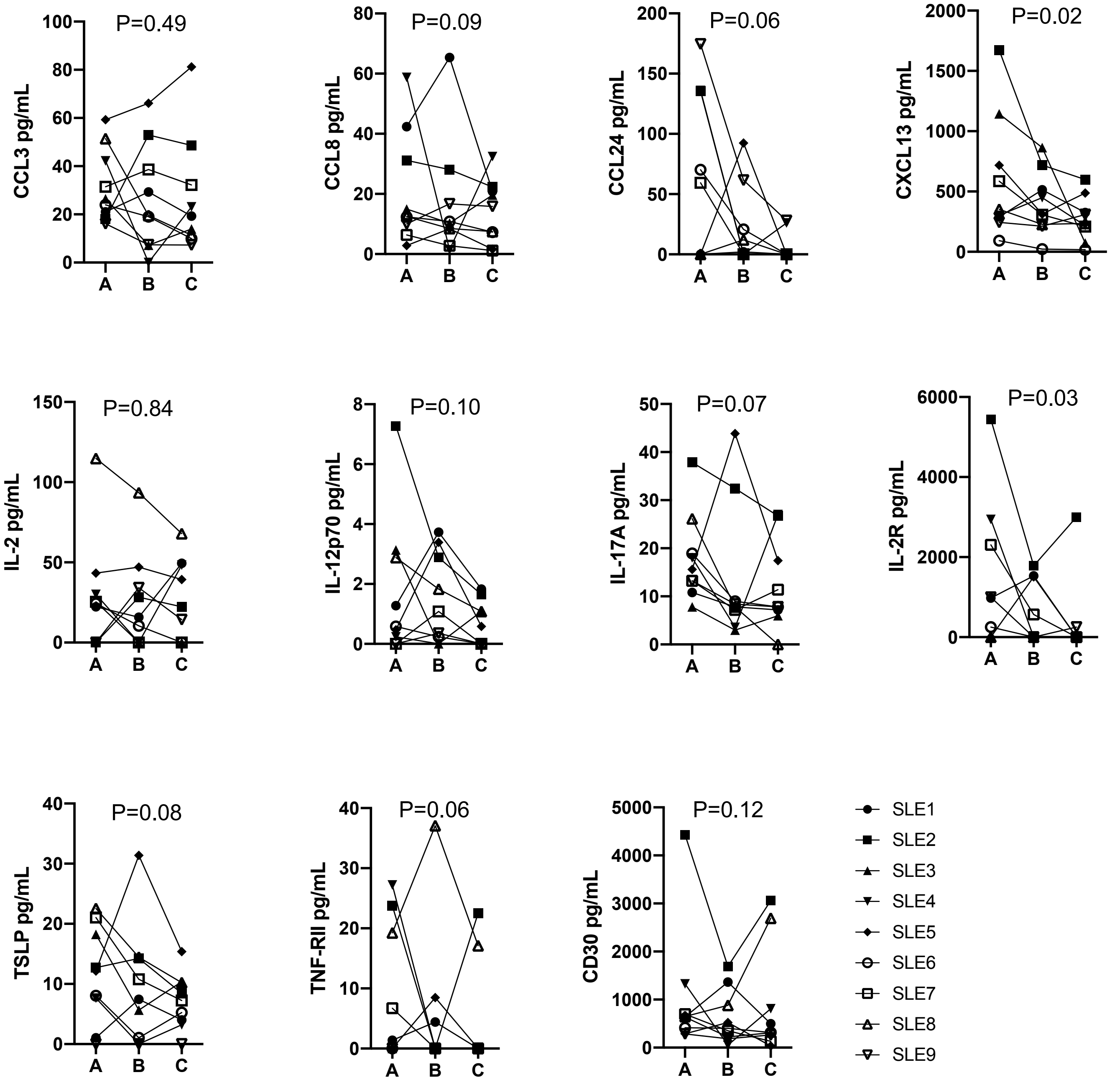
