## Supplementary figure legends for "Longitudinal immune cell profiling in early systemic lupus erythematosus"

**Supplementary Figure 1. Effect of treatment on CD19^int^ Ki67^hi^ plasmablasts in early SLE and established SLE patients**. (A) Correlation between prednisone dose and CD19^int^ Ki67^hi^ plasmablast frequency in nine early SLE and 15 established SLE. Spearman statistics shown. (B) Comparison of CD19^int^ Ki67^hi^ plasmablast frequency between the patients with immunosuppressive treatment (Prednisone>10mg and/or MMF) and those without. Nine early SLE and 15 established SLE were included in this analysis. Mann-Whitney U test was used for the comparison. *P<0.05, **P<0.01, ***P<0.001. Data are shown as mean ± SE

**Supplementary Figure 2. Longitudinal changes of CD19^int^ Ki67^hi^ plasmablasts and ABCs in early SLE.** Longitudinal changes of CD19^int^ Ki67^hi^ plasmablasts and ABCs in nine early SLE at enrollment (time A), six months after enrollment (time B), and 12 months after enrollment (time C). Wilcoxon matched-pair signed rank test to compare between A and C was used for calculation of P value.

**Supplementary Figure 3. Serum cytokine levels (18 cytokines) in controls and early SLE.** Comparison of serum cytokine levels between nine controls and nine early SLE (time A). Serum cytokines without significant differences between SLE patients and controls (18 cytokines) are shown. Mann-Whitney U test was used for the comparison. Data are shown as mean ± SE

**Supplementary Figure 4. Serum cytokine levels (11 cytokines) in controls and early SLE.** Comparison of serum cytokine levels between nine controls and nine early SLE (time A). Serum cytokines with significant differences between SLE patients and controls (11 cytokines) are shown. Mann-Whitney U test was used for the comparison. *P<0.05, **P<0.01, ***P<0.001. Data are shown as mean ± SE

**Supplementary Figure 5.** **Longitudinal evaluation of serum cytokine levels in early SLE patients.** Longitudinal changes of 11 cytokines in nine early SLE patients at enrollment (time A), six months after enrollment (time B), and 12 months after enrollment (time C). Wilcoxon log-rank test to compare between A and C was used for calculation of p-value.
